# DMT-induced shifts in criticality correlate with ego-dissolution

**DOI:** 10.1101/2025.02.08.636868

**Authors:** Mona Irrmischer, Marco Aqil, Lisa Luan, Tongyu Wang, Hessel Engelbregt, Robin Carhart-Harris, Klaus Linkenkaer-Hansen, Christopher Timmermann

**Affiliations:** GGZ Research, Academics Center for Trauma and Personality, Amsterdam, the Netherlands; DMT Research Group, Centre for Psychedelic Research, Imperial College London, London, United Kingdom; Department of Integrative Neurophysiology, Center for Neurogenomics and Cognitive Research (CNCR), Amsterdam Neuroscience, Vrije Universiteit Amsterdam, Amsterdam, The Netherlands; Weill Institute for Neurosciences, University of California San Francisco, San Francisco, CA, United States of America

**Keywords:** Psychedelics, Brain dynamics, DMT, EEG, Criticality

## Abstract

Psychedelics profoundly alter subjective experience and brain dynamics. Brain oscillations express signatures of near-critical dynamics, relevant for healthy function. Alterations in the proximity to criticality have been suggested to underlie the experiential and neurological effects of psychedelics. Here, we investigate the effects of a psychedelic substance (DMT) on the criticality of brain oscillations, and in relation to subjective experience. We find that DMT shifts the dynamics of brain oscillations away from criticality in alpha and adjacent frequency bands. In this context, entropy is increased while complexity is reduced. We find that the criticality shifts observed in alpha and theta bands correlate with the intensity ratings of ego-dissolution, a hallmark of psychedelic experience. Finally, using a recently developed metric, the functional excitatory-inhibitory ratio, we find that the DMT-induced criticality shift in brain oscillations is towards subcritical regimes. These findings have major implications for the understanding of psychedelic mechanisms of action in the human brain and for the neurological basis of altered states of consciousness.

## Introduction

Classic psychedelics elicit a wide range of changes in subjective experience and brain dynamics (Halberstadt, 2015; Nichols, 2016; Vollenweider and Preller, 2020; Timmermann et al., 2023a; Kelmendi et al., 2022; Aqil and Roseman, 2023). In recent years, thanks to a wealth of new and rigorous research, psychedelics have reemerged at the forefront of scientific and clinical inquiry due to their therapeutic potential and opportunities for research in brain function (Kyzar et al., 2017; Inserra et al., 2021; Bahji et al., 2023; Haikazian et al., 2023; van Elk and Yaden, 2022; Kwan et al., 2022).

The human brain has been suggested to operate in the proximity of a “critical point” (Bak et al., 1987; Cocchi et al., 2017). A prominent hypothesis concerning psychedelic states has been put forth by Carhart-Harris and colleagues (Carhart-Harris, 2018; Carhart-Harris et al., 2014; Carhart-Harris and Friston, 2019). The “entropic brain hypothesis” posits that 1) brain dynamics and subjective experience may be described in terms of their proximity to criticality; 2) the healthy, adult human brain operates in a slightly subcritical regime; and finally, 3) the effects of psychedelics may be characterized by a shift in criticality signatures, and in particular by an increase in entropy. Evidence in line with this hypothesis has been found by MEG, EEG, and fMRI studies (Pallavicini et al., 2021; Timmermann et al., 2023b, 2019; Schartner et al., 2017; Tagliazucchi et al., 2014; Atasoy et al., 2017; Toker et al., 2022; Varley et al., 2020; Ruffini et al., 2023).

Neuronal oscillations in the proximity of a critical point exhibit complex temporal patterns characterized by slowly decaying temporal autocorrelations of power-law form. These long-range temporal correlations (LRTC) represent memory of past activity in the signal and provide functional advantages for neuronal information processing (Avramiea et al., 2020, 2022; Bak, 1996; Shew and Plenz, 2013), likely contributing to the computational abilities of normal waking consciousness (Beggs, 2008; Bruining et al., 2020). The preservation of LRTCs is associated with healthy brain function (Beggs, 2008) and brain maturation is associated with increases in LRTC from childhood to early adulthood (Smit et al., 2011). Alterations in LRTC have also been found to be modulated by genetics (Linkenkaer-Hansen et al., 2007), sleep (Kantelhardt et al., 2015), sleep deprivation (Meisel et al., 2017)), anesthesia (Krzemiński et al., 2017), meditation (Irrmischer et al., 2018), and attention (Walter and Hinterberger, 2022).

Detrended fluctuation analysis (DFA) quantifies scale-free amplitude modulations of oscillations within a signal, thus revealing the presence of LRTCs, and proximity of brain oscillations to criticality (Hardstone et al., 2012; Linkenkaer-Hansen et al., 2001; Peng et al., 1995; Poil et al., 2012). DFA is also informative regarding the complexity and entropy of brain oscillations. In this context, entropy and complexity are inversely related. For example, a reduction in DFA estimates implies an increase in entropy, and a decrease in complexity, of the signal (and viceversa) (Rosso et al., 2007; Zanin and Olivares, 2021).

DFA provides information on distance from criticality of brain oscillations, but not on directionality (i.e., whether oscillations are supercritical or subcritical). A new measure was recently introduced for this purpose: the functional Excitatory/Inhibitory ratio (fE/I ratio) (Bruining et al., 2020). Unlike DFA, the fE/I ratio allows a distinction between excitation-dominated and inhibition-dominated regimes, thus distinguishing the directionality of criticality signatures observed with DFA (i.e., whether a given brain state operates in a subcritical or supercritical regime) (Bruining et al., 2020). DFA is still a prerequisite for the application of the fE/I ratio, which need not apply to oscillations that are too far from criticality.

In this work, for the first time, we quantify markers of criticality in two EEG-DMT datasets using DFA and the fE/I ratio, and compare the resulting changes with measures of subjective experience. We find 1) that DMT produces a shift *away* from criticality in alpha and adjacent frequency bands, 2) that the observed shift correlates with the ratings for the subjective experience of disruptions of the sense of self, and finally 3) that the observed shift is in the direction of subcritical regimes. Together, our findings provide novel insights into the mechanism of psychedelic action in the human brain and the neural correlates of self-related processing in normal waking and altered states of consciousness.

## Results

### DMT shifts dynamics of theta, alpha, and beta oscillations away from criticality; towards increased entropy and reduced complexity

In order to investigate the impact of DMT on the criticality of brain oscillations, we quantify the presence of LRTC following DMT and placebo administration in the same participants. We computed the DFA exponent as an index of LRTC. Paired-samples t-tests were conducted to compare the difference in DFA between placebo and DMT conditions for the theta, alpha and beta band. Compared to placebo, DMT induced a significant decrease of DFA in theta (ΔDFA_*θ*_ = −0.06; *p* = <.0001), alpha (ΔDFA_*α*_ = −0.09;*p* = <.0001) and beta (ΔDFA_*β*_ = −0.06; *p* = .0004). These observed reductions in DFA exponents were widespread throughout the scalp (Figure 2, left column), and are robust across subjects (Figure 2, left column boxplots). In sum, we find that DMT shifts the dynamics of brain oscillations in alpha and adjacent frequency bands *away* from criticality and in the direction of a more entropic, sub-critical regime. As implied above, DFA exponents also provide information on the entropy and complexity of brain signals. A DFA exponent of 1.0 represents a pink-noise-like signal (*1/f* spectrum), while a DFA exponent of 0.5 represents a white-noise-like signal (flat spectrum). Exponents with intermediate values represent the spectrum of fractional Gaussian noises between these two extrema. In this context, entropy and complexity are inversely related (Rosso et al., 2007; Zanin and Olivares, 2021). A white-noise-like signal has higher entropy but lower complexity, while a pink-noise-like signal has lower entropy but higher complexity. As such, a reduction in DFA exponent estimates, in the range between 0.5 and 1, such as that we observe here, indicates a shift away from a more complex, less entropic, pink-noise-like signal towards a less complex, more entropic, white-noise-like signal. In sum, our findings imply that brain oscillations in alpha and adjacent frequency bands under DMT become less complex, but more entropic, throughout the brain.

**Figure 1:**
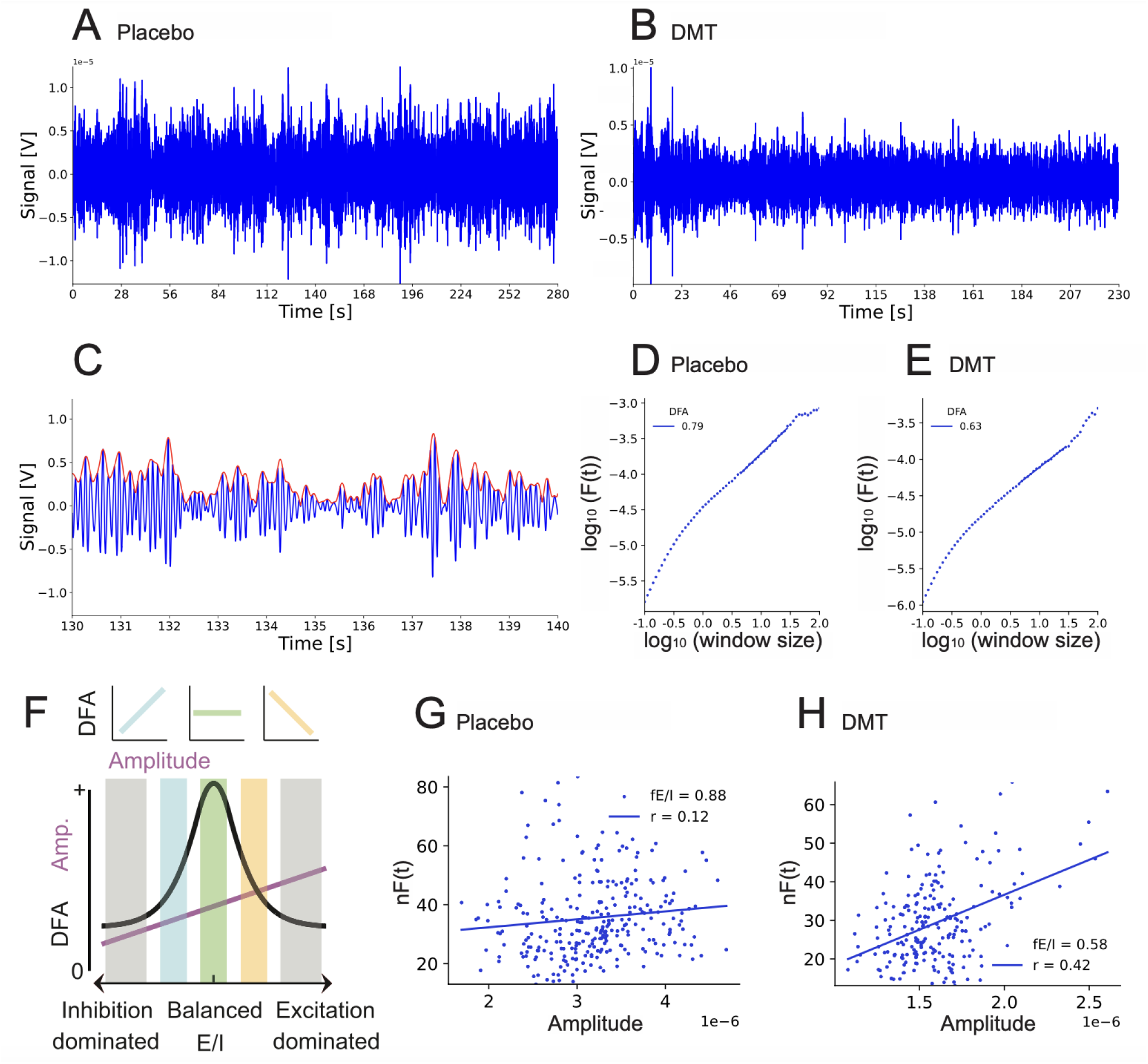
Assessing the criticality of neuronal oscillations under DMT and placebo. Examples shown of EEG from Pz filtered in the alpha-frequency range (8–13 Hz) during the placebo condition **(A)** and during DMT condition **(B)**. Note the change to more homogeneous variation in amplitudes during DMT compared to placebo (especially after approx. 20s, which corresponds to the onset of effects). **(C)** Amplitude of 10s alpha oscillations is shown in blue and the amplitude envelope in red. In detrended fluctuation analysis (DFA), amplitude modulations from placebo **(D)** and DMT **(E)** are plotted against window size in double-logarithmic coordinates to obtain DFA exponent, a marker for proximity to criticality. **(F)** The DFA exponent cannot distinguish sub-from super-critical activity, unless combined with the functional Excitatory/Inhibitory ratio (fE/I). fE/I of an example placebo signal **(G)** exhibits expected slight subcriticality close to the critical point, while the fE/I of DMT signal **(H)** shows a shift towards more subcriticality.

**Figure 2:**
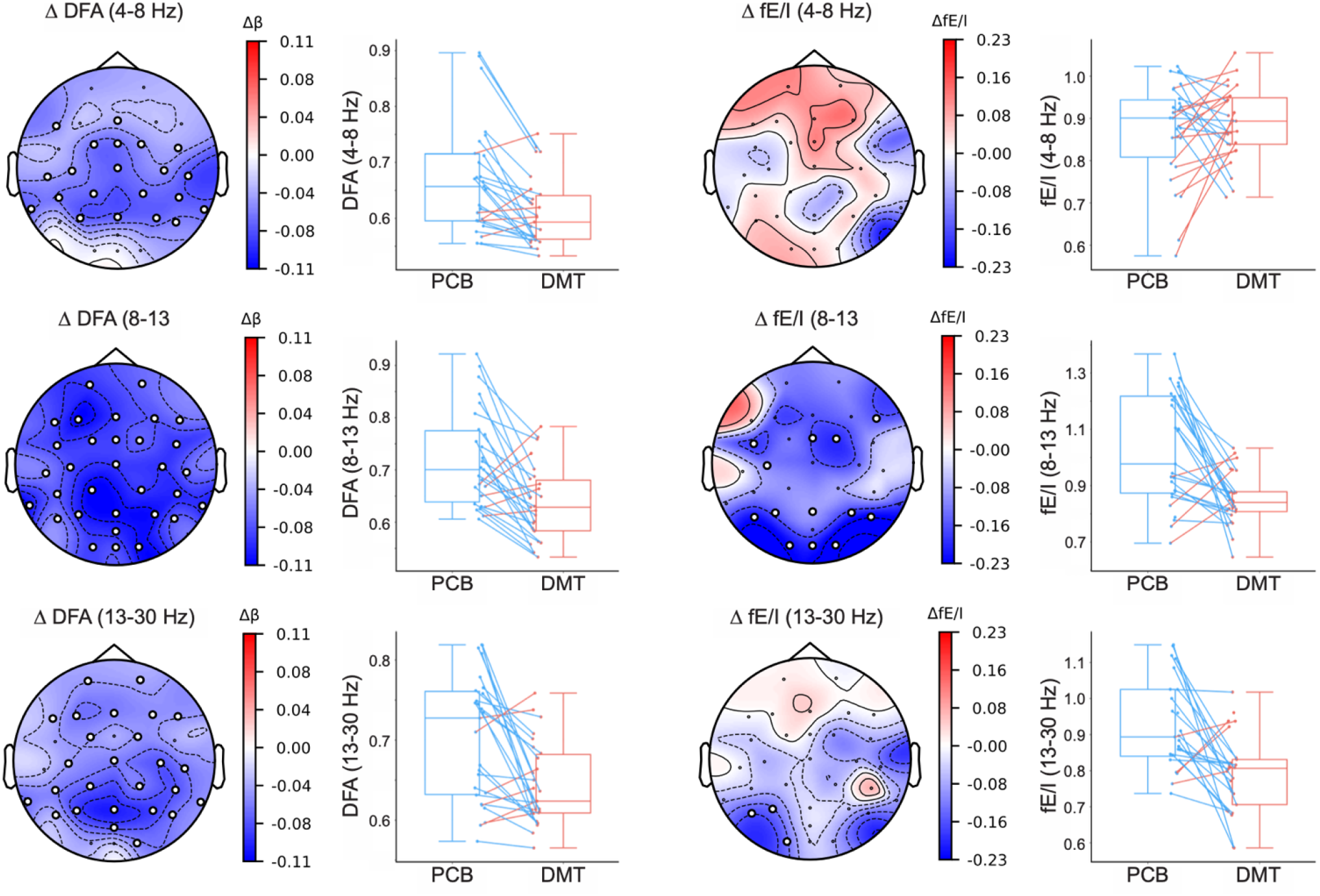
Brain oscillations under DMT in multiple frequency bands shift away from criticality, and towards (more entropic) subcritical regimes. Left, DFA exponents show statistically significant decreases in multiple frequency bands, throughout the scalp (electrodes with with circles). This indicates a shift away from critical dynamics in these frequency bands. Right, fE/I estimates show statistically significant decreases in the alpha band for occipital and parietal regions, and in the beta band for left-occipital region (electrodes with with circles). This indicates a shift towards more entropic, subcritical dynamics in these frequency bands and regions. Data points in box plots represent the mean of statistically significant electrodes per-participant.

### DMT shifts dynamics of alpha and beta oscillations towards subcritical regimes

DFA does not provide information on the direction of the criticality shifts it quantifies. Starting from a near-critical point, both shifts towards subcritical and supercritical dynamics are characterized by reductions in DFA exponents, such as those we observe here. To distinguish the directionality of the shift we observed, we computed a metric recently introduced for this purpose, the functional Excitatory/Inhibitory ratio (Bruining et al., 2020; Diachenko et al., 2024). Paired-samples t-tests were conducted to compare the difference in fE/I between the placebo and DMT condition. Compared to placebo, DMT induced a statistically significant decrease of fE/I for the alpha band (ΔfE/I_*α*_ = −0.18; *p* = .0003), which were especially pronounced in parietal and occipital electrodes. In the beta band the reduction only reached significance in the left occipital region (ΔfE/I_*β*_ = −0.14; *p* = .0005) (Figure 2, right column). In sum, we find evidence that the observed criticality shifts in alpha and theta bands are in the subcritical direction.

### Criticality shifts elicited by DMT correlate with the experience of ego-dissolution

Intense disruption of the sense of self - or ‘ego dissolution’ - is a hallmark of high-dose psychedelic experience, with potential relevance for therapeutic applications, and for understanding the neurological basis of conscious self-processing. To test whether shifts in criticality of neuronal oscillations measured with DFA and the fE/I ratio were associated with DMT-induced alterations in self-related processing, we performed correlations with 4 items of the Visual Analogue Scales (VAS) related to 3 elements related to disruptions in self-related processing as well as a generic item broadly assessing disruptions in the sense of self (see methods). We found statistically significant correlations between criticality shifts (measured with DFA estimates) and the generic item of self-disruption (measured with the item: *“I experienced a disintegration of my sense of self or ego”*) in the theta (*r*(25)=−0.61, *p*=.001) and alpha bands (*r*(25)=−0.56., *p*=.005). The correlation were statistically significant for most electrodes across the scalp (Figure 3). In sum, we find that the magnitude of the shift away from criticality elicited by DMT in theta and alpha bands relates to the disruptions in self-related processing.

**Figure 3:**
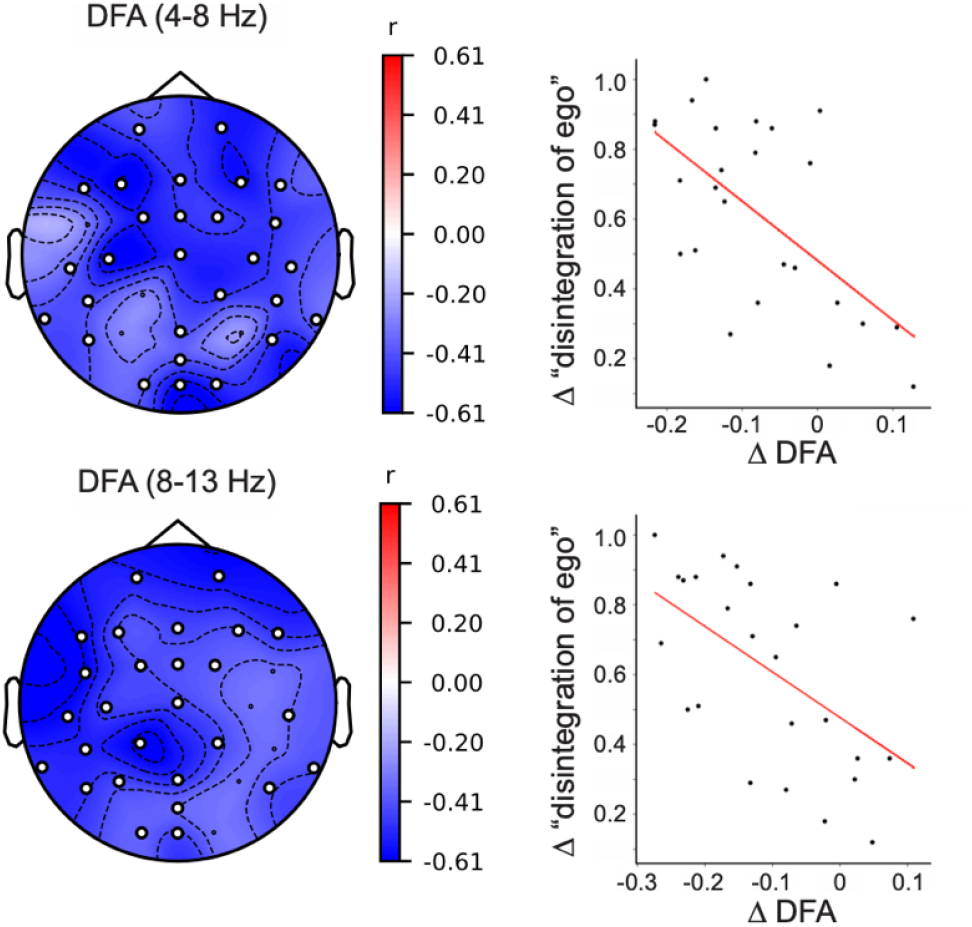
Criticality shift correlates with ego-dissolution experience. The change in DFA exponent in theta and alpha bands induced by DMT correlates with ego-dissolution experience. Left column, the correlation between ego-dissolution experience and DFA is statistically significant (filled circles) for electrodes throughout the scalp. Right column, correlation between mean DFA change in statistically significant electrodes of each participant and ratings of ego-dissolution.

## Discussion

Here, we investigated the effects of DMT, a classic psychedelic, on markers of criticality, and their relation to subjective experience. We first quantified changes in longrange temporal correlations (LRTCs) elicited by DMT using detrended fluctuation analysis (DFA) (Linkenkaer-Hansen et al., 2001; Poil et al., 2012). We found statistically significant decreases in the DFA exponent in theta, alpha, and beta bands throughout the scalp. This finding indicates a shift *away* from criticality for oscillations in these frequency bands, and implies a reduction in complexity and an increase in entropy. Next, we determined the directionality of the criticality shift, using a recently introduced metric, the functional Excitatory Inhibitory ratio (fE/I) (Bruining et al., 2020; Diachenko et al., 2024). We found statistically significant decreases of the fE/I ratio, indicating shifts towards subcritical regimes, in occipital and parietal alpha, and left-occipital beta bands. Finally, we correlated the observed changes in criticality markers with subjective measures of self-related processing, measured with Visual Anaolgue Scales (VAS). We found statistically significant correlations between DFA changes in alpha and theta bands and subjective intensity ratings of ego-dissolution experience, a hallmark of high-dose psychedelic experience. Our findings provide novel information on the effects of DMT on criticality (shifts *away* from critical point, and toward a more entropic, subcritical regime), i.e., entropy was increased, and complexity, decreased, in alpha and adjacent frequency bands. Furthermore, our findings demonstrate a relationship between criticality shifts and the disruption of the sense of self, providing novel candidates of neural correlates of self-related processing in the human brain.

Classic psychedelics are generally thought to be primarily active through the (excitatory) 5-HT2A receptor (Vollenweider and Preller, 2020); previous studies have reported increases in measures of signal diversity under psychedelics (Pallavicini et al., 2021; Timmermann et al., 2023b, 2019; Schartner et al., 2017); and weaker LRTCs are also observed in states of anesthetic unconsciousness and deep meditation, which certainly have profound differences with psychedelic states (Bruining et al., 2020; Irrmischer et al., 2018). Hence, our observation of weaker LRTCs and shifts towards subcritical (i.e. more inhibition-dominated) dynamics in alpha and adjacent frequency bands may appear puzzling at first. However, we believe that these apparently divergent findings can be reconciled. First, a key factor to consider is the multidimensional nature of conscious states. Certain states of consciousness (e.g. psychedelics and anesthesia) may be diametrically opposed in some dimensions (e.g. richness of content), while being similar in others (e.g. lack of a coherent sense of self). Indeed, we believe that the measures employed in our study selectively capture the shared nature of the latter similarity between otherwise distinct states of consciousness. As such, our study represents a significant step forward towards understanding the neural correlates of multidimensional consciousness.

A shift towards subcritical dynamics may intuitively be thought to indicate a decrease in entropy. However, this assumption is not always valid. A decrease in DFA exponent (between the range of 1 and 0.5) indicates a shift from a more pink-noise-like (DFA=1) process to a more entropic, white-noise-like (DFA=0.5) regime. The implication is that in a shift such as the one we observe, the entropy of the signal increases, while its complexity decreases (Rosso et al., 2007; Zanin and Olivares, 2021). The correlation with ego-dissolution indicates the functional relevance of this shift, which may be interpreted as a disruption of the habitual, structured and complex self-centered representations, which unfold over medium-long timescales, under the influence of DMT. On the other hand, our findings indicate that the suppression of complexity in alpha and adjacent frequency bands, indicated by weaker LRTCs and subcritical shifts, may hence provide the shared neural correlate between high-intensity psychedelic state and unconsciousness, corresponding to the loss of self as constitutional form of everyday experience. Indeed, previous work assessing LRTCs during deep meditation in experienced meditators also found a reduction in DFA exponent, correlating with absorption ratings (Irrmischer et al., 2018). Together, these findings provide concrete evidence of LRTCs as potential neural correlate for the habitual-waking sense of self, the stream of conscious thoughts present during regular waking consciousness, but absent or significantly disrupted during deep meditation, anesthesia, and intense psychedelic experiences. Alpha frequencies have been implied as top-down carriers of predictive models instantiated by high-level regions and networks (Alamia and VanRullen, 2019). Available theoretical frameworks and empirical evidence have indeed implicated reductions of alpha oscillations and default-mode network (DMN) connectivity (a system known to relate to alpha oscillations), as key players in psychedelic effects (Carhart-Harris and Friston, 2019). In fact, both systems have been found to be significantly dysregulated under DMT in an interrelated fashion (Timmermann et al., 2019). However, alpha oscillations also relate to activity in low-level sensory regions, which coexist in a dialogical interplay with high-level ones (Aqil and Roseman, 2023). Alpha oscillations and DMN activity are strongly suppressed by visual content, which is intensely present during DMT states (Timmermann et al., 2019). In fact, DMT-induced dysregulation of alpha oscillations has been found to significantly correlate with the intensity of visual experience (Timmermann et al., 2019). Recent empirical evidence has shown that even relatively simple simulated hallucinations can modulate highlevel cognition (Rastelli et al., 2022). Finally, we speculate that the cortical excitatory nature of DMT and other classic psychedelics (and the corresponding richness of conscious content) may be apparent in higher temporal frequency (high gamma), which are not accessible here because of experimental limitations. Changes of this nature have been observed in animal models of psychedelics (Yu et al., 2023). Alternatively, it is possible that theta oscillations may reflect supercritical regimes under DMT, as increases in theta have been found to be correlated with the visual experience induced by DMT (Timmermann et al., 2019). In this study, we found DFA to be reduced in theta oscillations; however, shifts in f(E/I) were inconclusive. Future studies should narrow the EEG analysis to moments of experience in which increases in content richness are present to advance these questions. This could be achieved by employing methodological paradigms that carefully match subjective experience with specific moments of the psychedelic state (Timmermann et al., 2023a).

In sum, we find that DMT alters the criticality signatures of brain oscillations, and that the observed shifts in criticality correlate with the subjective experience of ego-dissolution. In particular, we find that DMT shifts oscillations in alpha and adjacent frequency bands away from criticality and towards subcritical regimes characterized by increased entropy but reduced complexity. Weak, subcritical LRTCs in alpha and adjacent bands may be a potential shared neural correlate of the disruption of conscious self-related processing, common both to high-dose psychedelic experiences, deep meditation states, and anesthesia. Overall, our findings provide novel information on the effects of psychedelics on criticality, entropy, and complexity of brain oscillations, and their relation with subjective experience.

## Materials and Methods

### Participants and experimental procedures

For this study the EEG data from two placebo-controlled, single-blind, within-participant studies (Timmermann et al., 2019, 2023b) were combined resulting in 27 healthy participants (12 female, mean age= 34.1 SD=8.7) after removing participants with unusable EEG data due to muscle and movement artifacts. An initial screening visit to the Imperial College Clinical Research Facility (CRF) assessed physical and mental health to ensure suitability of participants. Exclusion criteria included: being under 18 years old, MR contraindications, no previous psychedelic experiences, an adverse reaction to a psychedelic, a history of psychiatric or physical illness rendering participants unsuitable for participation, a family history of psychotic disorder, or excessive use of alcohol or drugs of abuse. All participants provided written informed consent for participation in the study. Both studies were approved by the National Research Ethics Committee London—Brent and the Health Research Authority and was conducted under the guidelines of the revised Declaration of Helsinki (2000), the International Committee on Harmonization Good Clinical Practices guidelines, and the National Health Service Research Governance Framework. Imperial College London sponsored the research conducted under a Home Office license for research with Schedule 1 drugs.

Study 1 (12 participants from the final sample) was a dose-finding study where participants always received placebo (saline) on a first visit and DMT on a second visit one week after, while EEG recordings took place. The doses administered were as follows: 2 participants received 7 mg, 3 participants received 14 mg, one participant received 18 mg, and 4 participants received 20 mg of DMT fumarate - for full recruitment procedure and protocol see (Timmermann et al., 2019). In study 2 (15 participants from the final sample), participants received 20 mg of DMT fumarate and placebo (saline) in a counterbalanced order in visits separated 2 weeks apart - for full recruitment procedure and protocol see: (Timmermann et al., 2023b). For both studies, EEG recordings occurred at baseline (prior to administration) and for 20 minutes after intravenous administration of DMT fumarate, which was performed during 30 seconds and followed by a 15-second flush of saline. If participants partook in both studies, we only used the recordings corresponding to their participation in the second study to increase homogeneity in the doses. Participants attended the experimental sessions at the National Institute of Health Research (NIHR) Imperial Clinical Research Facility (CRF) for Study 1, and the Clinical Imaging Facility (CIF) at Imperial College London for Study 2.

For both studies, following EEG recordings and once subjective effects subsided, participants completed questionnaires and visual analogue scales designed to assess the subjective effects experienced during the administration of DMT (see (Timmermann et al., 2019, 2023b) for full procedures). Of these metrics, we selected 5 visual analogue scales for correlations with our EEG results, as these reflect central features of subjective experience we hypothesised are susceptible to changes in LRTC and changes in criticality: 1) the sense of self (‘I experienced a disintegration of my usual sense of ‘self’ or ‘ego’), 2) the sense of time (‘my sense of time was altered’), 3) the sense of space (‘my sense of size or space was altered’), 4) cognition (‘my thoughts wandered freely’). Additionally, we assessed how LRTC related to generic subjective effects induced by DMT (‘how intense was the drug experience’).

### EEG acquisition and preprocessing

The same EEG setup was used for both studies. Data were collected from 31 scalp locations in accordance with the 10-20 system using an MR-compatible BrainAmp MR amplifier (BrainProducts, Munich, Germany) and a compatible cap (BrainCap MR; BrainProducts GmbH, Munich, Germany). The system used FCz as the reference for all electrodes, and AFz was the ground electrode. Additionally, an ECG channel was used to capture heart rate recordings. For both recordings data were visually inspected, and segments of data containing muscle artifacts, head motion, and other gross artifact were removed prior to cleaning using Independent Component Analysis (see (Timmermann et al., 2019, 2023b) for a full list of preprocessing details).

### Quantifying long-range temporal correlations

The data were further analyzed with Neurophysiological Biomarker Toolbox (NBT) written in Python, github links available at (Diachenko et al., 2024). LRTC is a robust empirical feature of critical-state oscillations (Linkenkaer-Hansen et al., 2001, 2007; Monto et al., 2007). It estimates the temporal structure of oscillation amplitude and reflects the level of criticality in the network. Detrended fluctuation analysis (DFA) is used to assess LRTC in the signal in the time scales of interest via a DFA exponent. The DFA value of 0.5 indicates an uncorrelated random signal (i.e., absence of LRTC), whereas the value > 0.5 indicates the presence of positive auto-correlations and their strength. To quantify the strength of long-range temporal correlations (LRTC) in the amplitude modulation of the EEG oscillations, we first extracted the amplitude envelope using band-pass filters (FIR-filter, Blackman window with transition bandwidth of 1 Hz, (Figure 1A,B) and the Hilbert transform (Figure 1C). Next, the root-mean-square fluctuation of the integrated and linearly detrended signals, F(t), was calculated as a function of time window size, t (with an overlap of 50% between windows) and plotted in double-logarithmic coordinates (Figure 1D,E). The DFA exponent is the slope of the fluctuation function F(t) in a given interval, which was set to 2 to 30 seconds for the alpha band.

### Quantifying fE/I

The fE/I algorithm was developed based on an extended version of the Critical Oscillations (CROS) computational model of neuronal oscillations (Poil et al., 2012) which mimics the signals observed in human M/EEG recordings. In fE/I, the E/I ratio is estimated from the windowed covariation of the average amplitude and amplitude modulation (the temporal auto-correlation structure) of frequency-specific activity, in the presence of significant LRTC in the signal (Figure 1G,I). LRTC is a robust empirical feature of critical-state oscillations (Linkenkaer-Hansen et al., 2001, 2007; Monto et al., 2007) and is dependent on the E/I balance. It estimates the temporal structure of oscillation amplitude and reflects the level of criticality in the network. Detrended fluctuation analysis (DFA) is used to assess LRTC in the signal in the time scales of interest via a DFA exponent (Figure 1). The DFA value of 0.5 indicates an uncorrelated random signal (i.e., absence of LRTC), whereas the value > 0.5 indicates the presence of positive auto-correlations and their strength. Taken alone, however, the DFA exponent cannot distinguish sub-from super-critical activity (Figure 1F). Neither can the amplitude of oscillations, which changes monotonously with E/I balance, tell where the critical point is. The combination of the two, on the other hand, can be used to tell apart sub-, critical, and super-critical dynamics (Figure 1F). The correlation of amplitude and DFA is positive for a network operation in a slightly sub-critical state (Figure 1 F left), zero in a critical state (Figure 1F, middle), and negative in a slightly super-critical state (Figure 1F, right). Given that networks operating in these regimes exhibit co-variation in amplitude and temporal structure, we can use a sliding-window approach to quantify this covariation and, thus, infer the E/I balance of the underlying networks. Of note, when networks are far from criticality as reflected in DFA exponents < 0.6, there is no correlation between the co-variation of amplitude and temporal structure and, therefore, the DFA exponent of 0.6 is used as a threshold to compute the fE/I ratio (Bruining et al., 2020).

Given the presence of LRTC in the signal reflected by the DFA exponent greater than 0.6, fE/I is then computed by correlating the amplitude and LRTC in short windows. Windowed LRTC, in this case, is estimated through the normalized fluctuation function which serves as a reliable proxy of the DFA exponent on short time scales. Overall, the main steps of the fE/I algorithm are as follows (Bruining et al., 2020):

1. The signal is bandpass-filtered in the desired frequency range using a finite-impulse-response (FIR) filter.
2. The amplitude envelope of the filtered signal is extracted using Hilbert Transform.
3. The signal profile is calculated and segmented into 80%-overlapping 5-second windows.
4. The windows are normalized using the mean of the amplitude envelope calculated per window.
5. Subsequently, the normalized windows are detrended.
6. The normalized fluctuation function for each window is computed as the root-mean square fluctuation of the detrended amplitude-normalized signal profile.
7. Finally, the fE/I value is obtained by subtracting the Pearson correlation between the windowed normalized fluctuations and windowed amplitudes from 1. The fE/I is set to NaN (i.e., missing) if DFA exponent does not exceed the DFA threshold of 0.6.

### Statistical analyses

The EEG analysis was performed per channel with a non-directional paired t-test (level: *p* < 0.05), with average DFA across statistically significant electrodes reported. Changes in DFA and fE/I were calculated between the placebo and DMT condition to avoid order effects, while the correlations with changes in subjective experience was performed with changes from DMT baseline to DMT after injection, to minimize noise. Due to the continuous nature of subjective experience, the parametric Pearson’s correlation coefficient was used to test for correlations with experience. To prevent chance-level effects, we used the Benjamini Hochberg false discovery rate (FDR) multiple comparison correction method (Benjamini and Hochberg, 1995) as post hoc test with FDR adjusted q-values of < 0.10 within a specific frequency band.

## Acknowledgments

We would like to thank Marina Diachenko for her support with this project.

## CRediT authorship contribution statement

**Mona Irrmischer:** Conceptualization; Investigation; Software; Visualization; Writing - original draft; Writing - review & editing. **Marco Aqil:** Conceptualization; Investigation; Writing - original draft; Writing - review & editing. **Lisa Luan:** Conceptualization; Investigation; Writing - review & editing. **Tongyu Wang:** Conceptualization; Software. **Hessel Engelbregt:** Conceptualization; Supervision; Writing - review & editing. **Robin Carhart-Harris:** Conceptualization; Supervision; Writing - review & editing. **Klaus Linkenkaer-Hansen:** Conceptualization; Investigation; Software; Supervision; Writing - review & editing. **Christopher Timmermann:** Conceptualization; Investigation; Supervision; Writing - review & editing.

